# Degradation of *boscalid, pyraclostrobin, fenbuconazole*, and *glyphosate* residues by an advanced oxidative process utilizing ultraviolet light and hydrogen peroxide

**DOI:** 10.1101/2020.11.13.382440

**Authors:** Blake Skanes, Jordan Ho, Keith Warriner, Ryan S. Prosser

## Abstract

Recently an advanced oxidative process (AOP) combining H_2_O_2_ and UV-C light was observed to be effective at controlling *Listeria monocytogens* (Murray et al., 2018) and *Escherichia coli* O157:H7 and degrading chlorpyrifos residues on the surface of apples (Ho et al., 2020). Little is known about the application of AOP for the degradation of other pesticide residues. This study examined degradation of boscalid, pyraclostrobin, fenbuconazole and glyphosate by 3% (w/v) H_2_O_2_, UV-C (254 nm) irradiation and their combination on apple skin and glass. The extent of degradation was not significantly different between the AOP and optimal individual treatment. However, treatment susceptibility was different with glyphosate most effectively degraded by H_2_O_2_ exposure (up to 98% on apple, 3% (w/v) H_2_O_2_ at 30□C for 15 min) while boscalid, pyraclostrobin and fenbuconazole were more effectively degraded by UV-C (up to 88%, 100% and 70% degradation after ~11,000 mJ/cm^2^). Suggestions for possible causes of degradation are proposed.

## 1 Introduction

Around the world an estimated 2.5-3 billion kilograms of pesticide are applied annually (Alavanja, 2009; Pimentel & Burgess, 2014). Agricultural use of pesticides remains a convenient method to meet global food demand and agricultural use makes up the largest portion of annual pesticide use in the world (Baylis, 2000; Pimentel & Burgess, 2014; Woodburn, 2000). Glyphosate is the most widely used pesticide in the world and has recently been the centre of controversy following debate regarding its carcinogenicity between the International Agency for Research on Cancer and many national governments (EPA, 2019; IARC, 2017; Tarazona et al., 2017). While glyphosate is broadly used in agriculture for weed control, fungicides are critically important in agriculture for food safety and to minimize spoilage (Gianessi & Reigner, 2006; Moss, 2008). While most pesticides have a pre-harvest interval (PHI), a minimum window of time required between the final application of the pesticide and the date of harvest, with some produce fungicides do not have a PHI (DOW AgroSciences, 2011; BASF, 2015). While there are still maximum residue limits (MRLs) that must be adhered to for these food commodities, the absence of a PHI in some instances makes fungicides important targets for post-harvest techniques that can degrade fungicide residues.

Boscalid, pyraclostrobin and fenbuconazole are fungicides commonly used in many agricultural applications in Canada, they are approved for use on 318, 417, and 58 food commodities respectively (PMRA, 2020) (Fig. 1). In Canada, boscalid and pyraclostrobin are the active ingredients in the product PRISTINE® and this product does not have a PHI for field berries, mechanically harvested cucurbit vegetables, carrot, celeriac, greenhouse cucumbers, lettuce and tomatoes, celery, spinach, and stone fruit. Fenbuconazole is the active ingredient in the product INDAR® and it does not have a PHI for citrus fruits (DOW AgroSciences, 2011; BASF, 2015).

**Figure 1:**
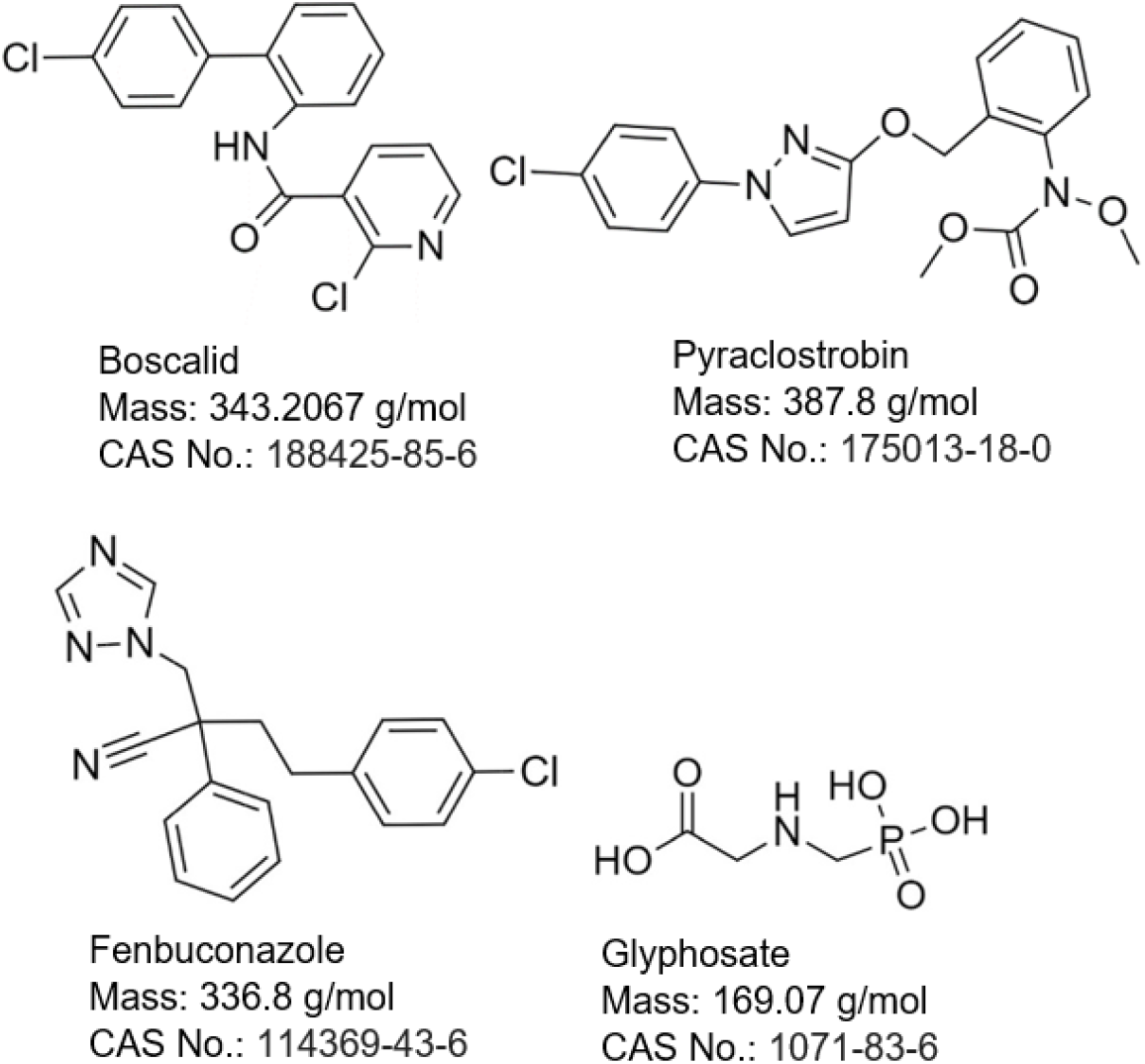
Chemical structure of boscalid, pyraclostrobin, fenbuconazole and glyphosate.

While the Canadian Food Inspection Agency’s (CFIA) National Chemical Residue Monitoring Program consistently reports that maximum residue limit (MRL) compliance rates for all tested commodities are >95%, public perception is that pesticide residues on food are a risk to human health. The majority of samples that are non-compliant are technical violations of the 0.1 μg/g default MRL for commodity/pesticide combinations for which Health Canada has not yet established an MRL (CFIA, 2015; 2017). Of particular concern to consumers are the twelve commodities, termed the dirty dozen, that are most often found to have multiple detectable pesticide residues on their surface (Lantz, 2015). Apples are found at the top of the dirty dozen list and regulatory testing has shown that 73% of Canadian domestic and 88% of imported apples test positive for multiple surface residues on their way to market (it is important to note the distinction that these are detected residues, not residues exceeding the MRL) (CFIA, 2015). With apples making up 20% of Canada’s $1 billion fruit industry, there is commercial interest in techniques that can help with residue limit compliance and the public’s concern about pesticide residues on apples (StatsCan, 2018).

Pesticide degradation using a number of common techniques is a well-studied field. Washings with water, chlorine baths, ozonated water, ozone, mild acids and hydrogen peroxide have been documented to be somewhat effective (summarized in Table S1 of the SI) at the removal of pesticide residues from the surface of fruits and vegetables (Hoff et al., 2019; Lester et al., 2017; Sadło et al., 2017; Savi et al., 2016; Bajwa & Sandhu, 2014; Cengiz & Certel, 2014; Junges et al., 2013; Karaca et al., 2007; 2012; Kusvuran et al., 2012; Wu et al., 2009; 2007; Crowe et al., 2006; Ikehata & Gamal El-Din, 2005; Hwang et al., 2001b; 2002; 2003). Using these wash techniques can increase the risk of microbial contamination (CDC, 1989; DiCaprio et al., 2015; Franz et al., 2008; Gombas, 2017). A potential alternative is the use of advanced oxidative processes (AOPs), for example the combinations of multiple treatments such as UV light, hydrogen peroxide, ozone, titanium dioxide and others that are capable of generating hydroxyl or other highly reactive radicals that can degrade pesticide residues (Junges et al, 2013; Lassalle et al., 2014; 2015; Manassero et al, 2010). AOP has been a common technique to degrade pollutants in wastewater for decades and has recently seen some application in the treatment of food commodities for the same purpose. Recently, an AOP was developed in the Department of Food Science at the University of Guelph for the treatment of apples, and other produce, in response to a *Listeria monocytogenes* outbreak (Glass et al., 2015). The process combines ultraviolet light (254 nm generated by low pressure mercury lamps) and hydrogen peroxide (6%, applied in a spray at 48□C), and has been shown to reduce *L. monocytogens* (3 log CFU reduction) and *Escherichia coli* O157: H7 (>6.6 log CFU reduction) on apples (Murray et al., 2018; Ho et al., 2020). Since AOPs enable biological control through interaction with organic compounds to cause bacterial and fungal cell death, they hold potential for the degradation of pesticide residues as well.

The apple skin presents a relevant medium for the study of pesticide residue degradation by AOP. Owing to the waxy cuticle pesticide residues typically remain on their surface following application (Angioni, et al., 2004). On the surface the pesticide residues can receive a greater exposure to treatment conditions compared to other types of produce, where the residue may be absorbed. Boscalid, pyraclostrobin and fenbuconazole were selected as fungicides with wide applicability, and relevance to apple production while glyphosate was selected as being of general interest as a target for residue degradation considering its wide use in agriculture (IARC, 2017). This study aimed to study the degradation of the residues of these four pesticides after treatment with hydrogen peroxide, UV light or an AOP combining these treatments on both apple skin and glass. The objective of the study was to determine whether the AOP is capable of significant degradation of pesticide residues in addition to the reduction of pathogenic bacteria it has demonstrated.

## 2 Materials and Methods

### 2.1 Chemicals

Glyphosate, fenbuconazole and boscalid (≥98% purity) PESTANAL analytical standards were obtained from Sigma Aldrich (Oakville, ON, Canada). Pyraclostrobin was obtained from Toronto Research Chemicals (Toronto, ON, Canada). HPLC grade acetonitrile (99.9% purity) and methanol (99.8% purity) were obtained from Caledon Laboratory Chemicals (Georgetown, ON, Canada). Acetone (99.5% purity, certified ACS) was obtained from Fisher Scientific (Ottawa, ON, Canada). Hydrogen peroxide was obtained from a local grocery store and was 3% w/v (0.88 mol/L). Water used in the preparation of glyphosate solutions was distilled in the laboratory. Apples (*Malus domestica* Mutsu, aka Crispin) were obtained directly from a local farmer (Niagara, ON, Canada).

### 2.2 Spike Solutions and Application/Recovery of the Pesticides

To keep matrix effects to a minimum, solvent rinses were used for recovery rather than popular whole fruit methods like QuEChERS that require tissue homogenization. Methanol, acetonitrile, and acetone were tested as application and recovery solvents. For the fungicides, application with acetonitrile and recovery with methanol proved to be the most effective with recoveries up to 94%, 96% and 99% on glass for boscalid, pyraclostrobin and fenbuconazole respectively (87%, 80% and 70% respectively on apple skin). Glyphosate, owing to its poor solubility in the tested solvents, was instead applied and recovered with distilled water (up to 100% and 96% recovery on glass and apple respectively). Data on the performance of the different solvents for recovery are provided in Tables S2 and S3 of the SI.

The treatment of apple and glass samples prior to exposure to the various oxidative treatments varied between the fungicides and glyphosate. Pesticides were applied either directly to a small glass petri dish or ~2 cm × 2cm segment of apple skin laid flat on a small petri dish. To prevent the application solvent from interfering with the pesticide’s exposure to the oxidative treatment, the application solvent was allowed to evaporate completely prior to exposure. These solvents would not be present when a pesticide is applied to an apple in the field. When the apple is harvested, the droplets of pesticides will have dried after the last application. A 50-μL aliquot of spike solution (5.827mmol/L (2,000 μg/L) boscalid, 25.785 mmol/L (10,000 μg/L) pyraclostrobin, 29.691 mmol/L (10,000 μg/L) fenbuconazole and 59.147 mmol/L (10,000 μg/L) glyphosate) was pipetted onto glass or apple skin and allowed to dry completely in the dark. The 50-μL aliquot corresponds to the average volume of a single droplet of water and would thus represent the largest volume of pesticide that would likely coalesce during spray application without dripping off the fruit (and thus the largest concentration). The pesticides applied in acetonitrile took between 10 and 15 minutes to dry depending on laboratory conditions. Glyphosate samples were dried in a 50□C oven for 15 to 30 minutes, as at room temperature spikes took up to 3 hours to dry. Following exposures, 1mL of recovery solvent (methanol for fungicides, distilled water for glyphosate) was pipetted onto the application site. For recovery from apple skin the dish was held at an angle to allow the solvent to collect at the periphery, limiting absorption into the exposed flesh, the segment was flipped over and the dish was returned to horizontal to allow the solvent to contact the skin for several seconds with gentle swirling after which the solvent was collected in LC vials. Samples on glass were recovered in the same fashion. The nominal concentrations of the recovered pesticides were 0.291 mmol/L (100 μg/L) for boscalid, 1.289 mmol/L (500 μg/L) for pyraclostrobin, 1.485 mmol/L (500 μg/L) for fenbuconazole and 2.957 mmol/L (500 μg/L) for glyphosate.

Positive controls were spiked with pesticide and not treated with the oxidative treatments; effectively serving as a recovery control. All exposures included positive controls. Negative controls were not spiked but were exposed to the oxidative treatments. Negative controls were used for ultraviolet light and AOP exposures. During exposures involving hydrogen peroxide, the positive control involved the spike with pesticides and an additional 100-μL of recovery solvent (methanol for the fungicides, distilled water for glyphosate) while the negative controls were not spiked and received 100-μL of 3% (w/v) hydrogen peroxide.

### 2.3 Hydrogen Peroxide Exposures

Following the application of the pesticides 100 μL of 3% (w/v) hydrogen peroxide was pipetted onto glass or apple skin in the location of the applied pesticide. The 100-μL aliquot provided 0.088 mmol of hydrogen peroxide which had previously been shown to minimize scavenging of the hydroxyl radical by hydrogen peroxide (Glaze et al., 1995) and is within the typical range for use as a bactericide (Parish et al., 2003). A hot water bath was used to control the temperature of the hydrogen peroxide at the time of application. Room temperature (25 ± 3□C), 30□C, 40□C, 50□C and 60□C were tested. Exposures to hydrogen peroxide lasted for 15 minutes at room temperature. Samples were then analyzed by liquid chromatography coupled with mass spectrometry.

### 2.4 UV Exposures

Following the spiking of glass or apple, samples were loaded onto a plastic tray and placed under a hood containing 4 × 25 W lamps (Sani-ray, Happauge, NY,USA) emitting UV light at 254 nm. Incident UV light intensity (mW/cm^2^) was measured between exposures. Samples were maintained at the same height as the radiometer (Trojan Technologies Inc, London, ON, CA) to ensure that the exposure of the samples to UV was measured accurately. Exposure durations were 30 s, 1, 2, 5, 10, 15, 30, or 40 min. Exposure times were converted to UV dose (mJ/cm^2^) by multiplying the intensity of the UV light by the exposure duration. Often the UV intensity decreased the longer the bulbs were in operation (for example the initial intensity of 5.71 mW/cm^2^ decreased to 4.89 mW/cm^2^ immediately after the 40 min exposure). The decrease was assumed to be linear and the dose was adjusted according to the initial and concluding intensity (i.e. the average of the UV intensity measured prior to insertion of the sample and after removal was used to determine the dose).

### 2.5 Advanced Oxidative Process

Following sample preparation, as outlined above, 100 μL of 3% (w/v) hydrogen peroxide at the optimal application temperature, as determined in the hydrogen peroxide exposures (room temperature for fenbuconazole and glyphosate, 60□C for boscalid and pyraclostrobin), was pipetted onto the sample immediately prior to sample placement under the UV lamps. Exposure times were the same as those used for the UV exposures (thus UV dose was comparable) and were therefore different than those for the hydrogen peroxide treatments. UV intensity was monitored at the initiation and conclusion of exposure and was converted to UV dose (mJ/cm^2^) using exposure duration. As with the hydrogen peroxide exposures, positive controls were spiked with an additional 100 μL aliquot of recovery solvent prior to recovery.

### 2.6 Chemical Analysis

Liquid chromatography-mass spectrometry analyses were performed in the Mass Spectrometry Facility within the Advanced Analysis Centre at the University of Guelph. Samples were injected into a Dionex UHPLC UltiMate 3000 liquid chromatograph interfaced to an amaZon SL ion trap mass spectrometer (Bruker Daltonics, Billerica, MA, USA). An Eclipse Plus C18 column (5-micron particle size, 150 mm × 2.1 mm, Agilent, Santa Clara, CA, USA) was used for chromatographic separation. While all chromatography methods utilized water (0.1% formic acid) and acetonitrile (0.1% formic acid), conditions were optimized for each pesticide (Table S4). The flow rate was maintained at 0.4 mL/min. The mass spectrometer electrospray capillary voltage was maintained at 4.5 kV with a drying gas temperature of 300 C with a flow rate of 12 L/min. Nebulizer pressure was 40 psi. Nitrogen was used as both nebulizing and drying gas while helium was used as collision gas at 60 psi. The mass spectrometer was set on Single Reaction Monitoring (SRM) for all pesticides. Ion mode and the monitored transition for each pesticide is presented in the SI (Table S5). Detection and quantification limits were based on low concentration standards and corresponded to the concentrations producing signal to noise ratios of 3 and 10 respectively, or slightly greater if the next lowest concentration dropped below that threshold. Boscalid had an LOD 0.5 μg/L and LOQ 1 μg/L. Pyraclostrobin and Fenbuconazole both had LOD and LOQ < 0.5 μg/L. Glyphosate had LOD 15 μg/L and LOQ 31.25 μg/L.

### 2.7 Statistical Analysis

One-way analysis of variance (ANOVA) (α = 0.05) was used to compare the degradation of pesticides among treatments. RStudio (version 3.6.2) was used toe perform ANOVA. The Shapiro-Wilk and Bartlett tests were used to determine normality and equal variance among treatments. A Tukey’s HSD test was performed to determine significant pairwise difference between treatments. If the normality test failed, square root, cube root and log transformations of the data were attempted. If a transformation successfully achieved normality, ANOVA was conducted on the adjusted data. If transformation failed to achieve normality or the equal variance test failed, a Kruskal-Wallis test on ranks was completed to test for significant differences among treatments followed by a Dunn’s multiple comparison test.

## 3 Results and Discussion

### 3.1 Boscalid

While boscalid was not significantly degraded on glass when 3% hydrogen peroxide was applied at 30□C (Fig. 2), degradation was significant at 40□C, 50□C and 60□C (Table 1) with the greatest residue degradation occurring at 60□C (0.106 ± 0.007 mmol degradation, ~38%) (Table 2). On apple skin the boscalid control was poorly recovered resulting in a significant difference compared to all treatment temperatures (Fig. 3). It should be noted that while the boscalid control on apple skin was poorly recovered variability in the recovered boscalid for each exposure was relatively small. While it is thus not possible to estimate the amount of boscalid that was degraded at each temperature, the recovered quantity was again the lowest at 60□C compared with the other treatment temperatures which suggests greater degradation. No previous studies examining the degradation of boscalid by hydrogen peroxide have been reported in the literature.

**Figure 2:**
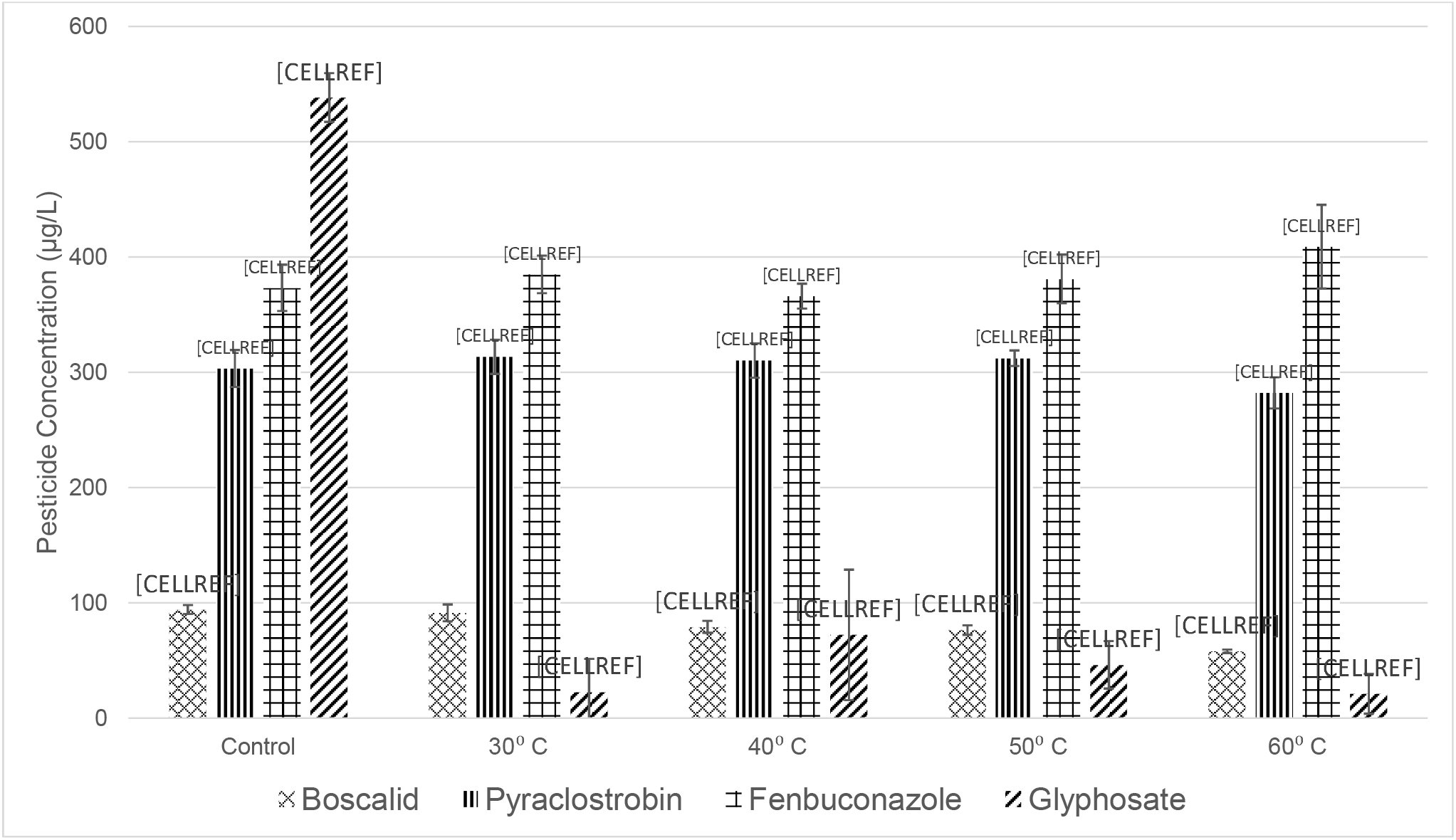
Recovered concentration of boscalid, pyraclostrobin, fenbuconazole and glyphosate following exposure to 3% w/v hydrogen peroxide for 15 minutes at various temperatures on glass (n = 3). Temperature is representative of temperature at time of application. Error bars represent a standard deviation. * Denotes a significant difference from the control.

**Figure 3:**
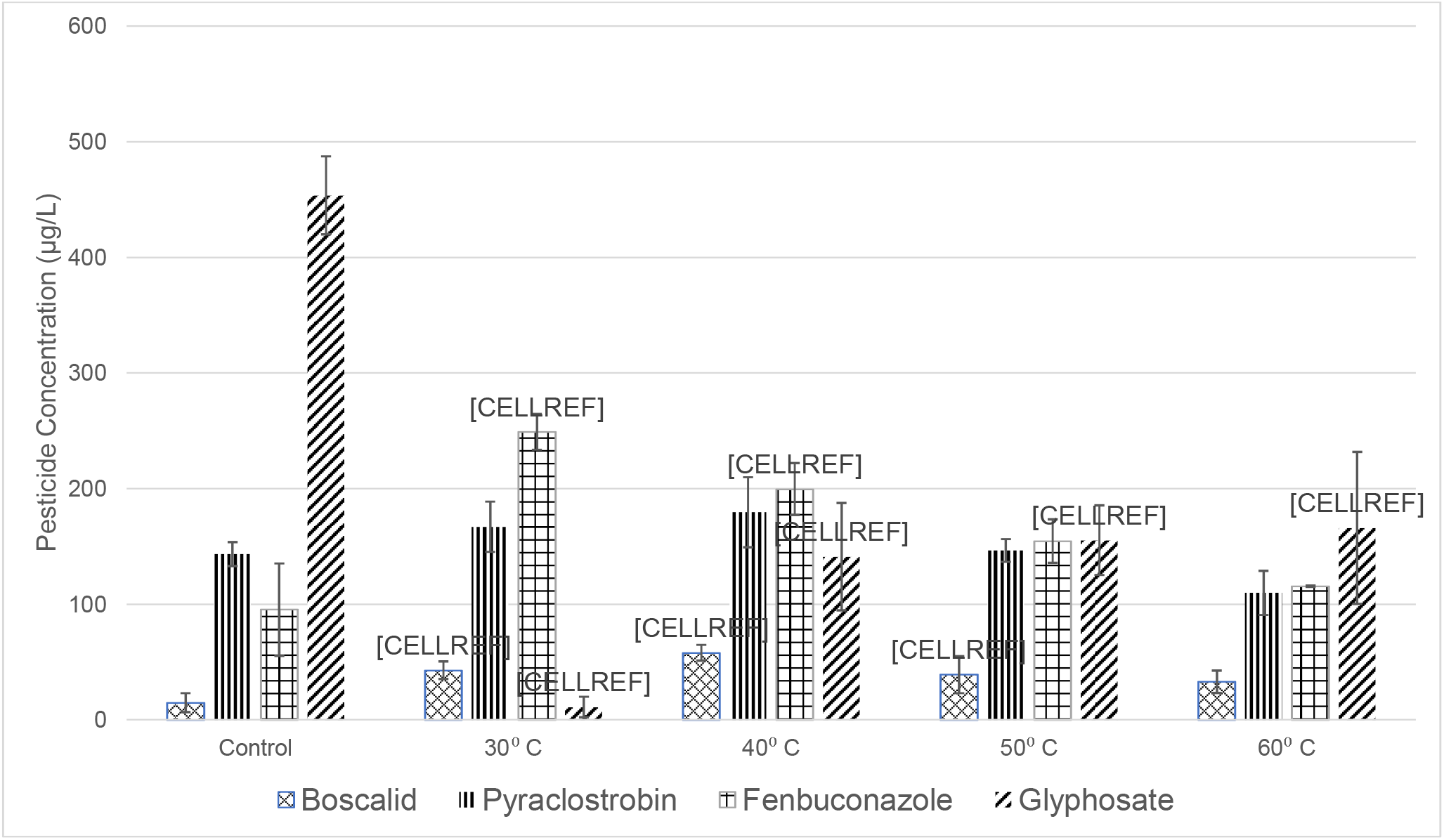
Recovered concentration of boscalid, pyraclostrobin, fenbuconazole and glyphosate following exposure to 3% w/v hydrogen peroxide for 15 minutes at various temperatures on apple skin (n = 3). Temperature is representative of temperature at time of application. Error bars represent a standard deviation. * Denotes a significant difference from the control.

**Table 1:** Normalized recoveries of boscalid, pyraclostrobin, fenbuconazole and glyphosate after treatment on glass (top) or apple skin (bottom) with 3% w/v hydrogen peroxide. N=3. Data was normalized with respect to the positive controls. Superscripts denote a significant difference between labelled exposure and ^a^ control, ^b^ 30□C, ^c^ 40□C.

| Compound | Control | 30 °C | 40 °C | 50 °C | 60 °C |
| --- | --- | --- | --- | --- | --- |
| Glass |  |  |  |  |  |
| Boscalid | 1 | 0.969 | 0.840<sup>a</sup> | 0.812<sup>ab</sup> | 0.616<sup>abc</sup> |
| Pyraclostrobin | 1 | 1.033 | 1.023 | 1.029 | 0.930 |
| Fenbuconazole | 1 | 1.031 | 0.981 | 1.021 | 1.096 |
| Glyphosate | 1 | 0.043<sup>a</sup> | 0.135<sup>a</sup> | 0.086<sup>a</sup> | 0.040<sup>a</sup> |
| Apple |  |  |  |  |  |
| Boscalid | 1 | 2.863<sup>a</sup> | 3.871<sup>a</sup> | 2.612<sup>a</sup> | 2.202 |
| Pyraclostrobin | 1 | 1.165 | 1.252 | 1.021 | 0.767<sup>c</sup> |
| Fenbuconazole | 1 | 2.608<sup>a</sup> | 2.090<sup>a</sup> | 1.620<sup>b</sup> | 1.211<sup>bc</sup> |
| Glyphosate | 1 | 0.025<sup>a</sup> | 0.311<sup>ab</sup> | 0.343<sup>ab</sup> | 0.366<sup>ab</sup> |

**Table 2:** Molar degradation of boscalid, pyraclostrobin, fenbuconazole and glyphosate after treatment on glass with 3% w/v hydrogen peroxide (n = 3).

|  | **Boscalid** |  | **Pyraclostrobin** |  | **Fenbuconazole** |  | **Glyphosate** |  |
| --- | --- | --- | --- | --- | --- | --- | --- | --- |
|  | Num. Moles degraded from control (mmol) | St Dev | Num. Moles degraded from control (mmol) | St Dev | Num. Moles degraded from control (mmol) | St Dev | Num. Moles degraded from control (mmol) | St Dev |
| **Control** | 0 | ±0.009 | 0 | ±0.034 | 0 | ±0.049 | 0 | ±0.102 |
| **30□C** | 0.008 | ±0.014 | -0.026 | ±0.033 | -0.035 | ±0.044 | 3.050 | ±0.122 |
| **40□C** | 0.044 | ±0.011 | -0.018 | ±0.033 | 0.021 | ±0.039 | 2.757 | ±0.206 |
| **50□C** | 0.052 | ±0.010 | -0.023 | ±0.026 | -0.023 | ±0.050 | 2.911 | ±0.101 |
| **60□C** | 0.106 | ±0.007 | 0.055 | ±0.031 | -0.106 | ±0.071 | 3.060 | ±0.093 |

Boscalid was found to be susceptible to UV light on both glass (Fig. 4) and apple skin (Fig. 5) with significant degradation occurring within 30 s (150 mJ/cm^2^) on glass and 10 min (2985 mJ/cm^2^) on apple skin (Tables 3 & 4). On glass, there was a significantly greater degradation of boscalid residues (0.217 ± 0.010 mmol) after exposure to a 11040 mJ/cm^2^ UV dose (40 min) compared to all other exposures whereas on apple skin there was no significant difference in degradation after 2985 mJ/cm^2^ (10 min) (Tables 3 & 4). This corresponded to ~88% and up to 100% degradation on glass and apple skin, respectively. Lassalle et al. (2014) have previously observed that UV irradiation effectively degrades boscalid residues using a high-pressure mercury lamp irradiating 120 mL quartz tubes in a sonicator at 25 ± 3□C. While the UV dose to samples was not reported, 150 min of exposure of a 5 mg/L aqueous solution of boscalid to a lamp with a radiation flux of 6200 lumens (wavelength 200 to 650 nm) resulted in a 90% decrease in residues. More rapid degradation of boscalid was observed in the current study. This may be due to the water in the quartz tubes used by Lassalle et al. (2014) reducing the intensity of the incident UV light compared to that at the bulb, the difference in the wavelength of light produced by the bulbs in each study (254 nm vs 200 to 650 nm) or the formation of reactive oxygen species in the current study through photoelectron transfer of the absorbed energy to oxygen in the air surrounding the exposure (see section 3.5 for more detail on photoelectron transfer). Lagunas-Allué et al. (2010) did not observe significant degradation (8%) of boscalid when 25 mL of a 3.5 mg/L solution of boscalid was irradiated with 365 nm UV light for 90 min. As both Lagunas-Allué et al. (2010) and Lassalle et al. (2014) used an aqueous solution for their exposures it is likely that the lack of significant decrease in residue observed by Lagunas-Allué et al.(2010) is due to the wavelength of UV used. The wavelength of UV light used for boscalid residue degradation should therefore be carefully selected when used without a catalyst.

**Figure 4:**
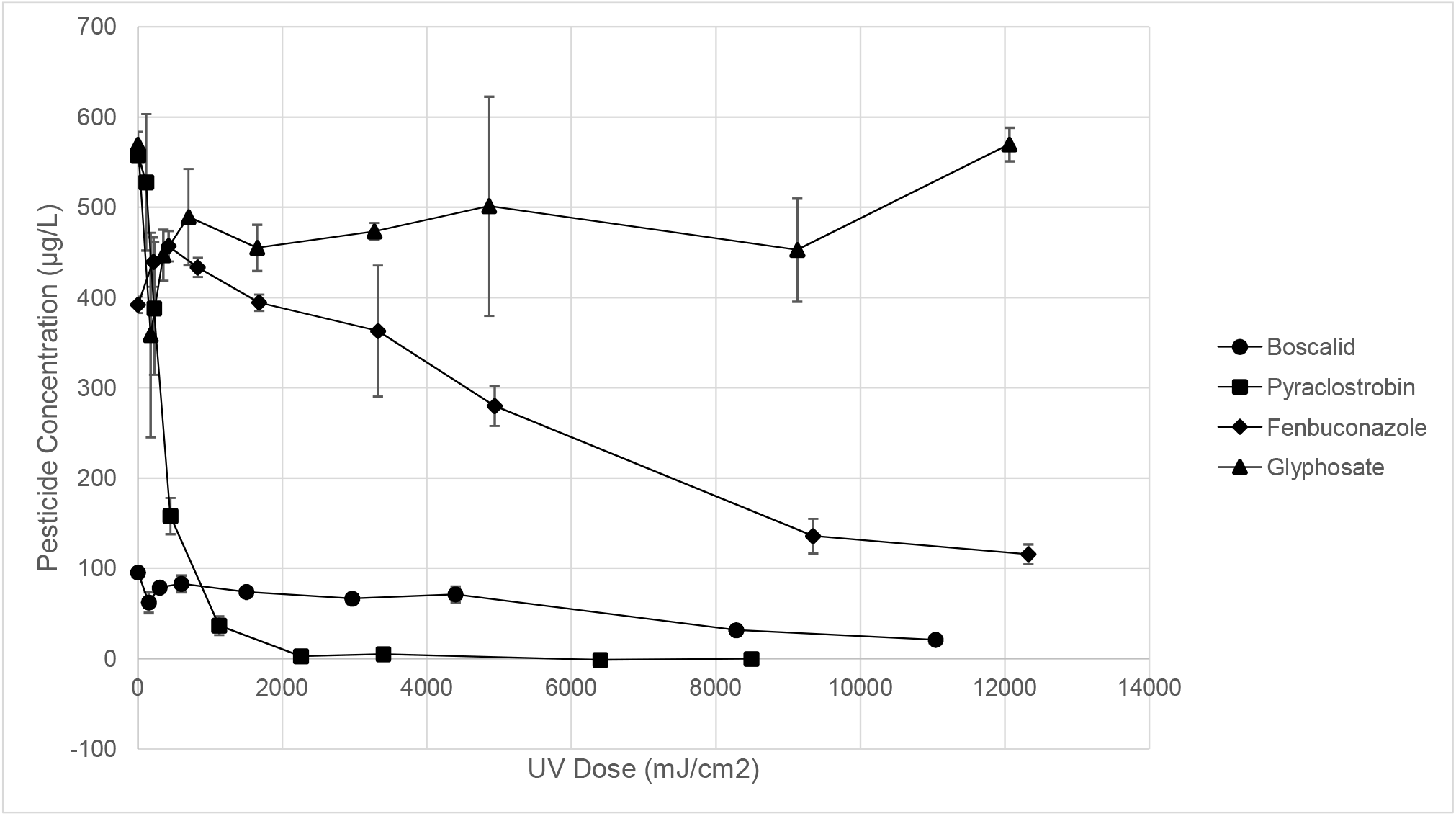
Recovered concentrations of boscalid, pyraclostrobin, fenbuconazole and glyphosate following exposure to 254 nm UV light at various UV doses on glass (n = 3). Error bars represent a standard deviation.

**Figure 5:**
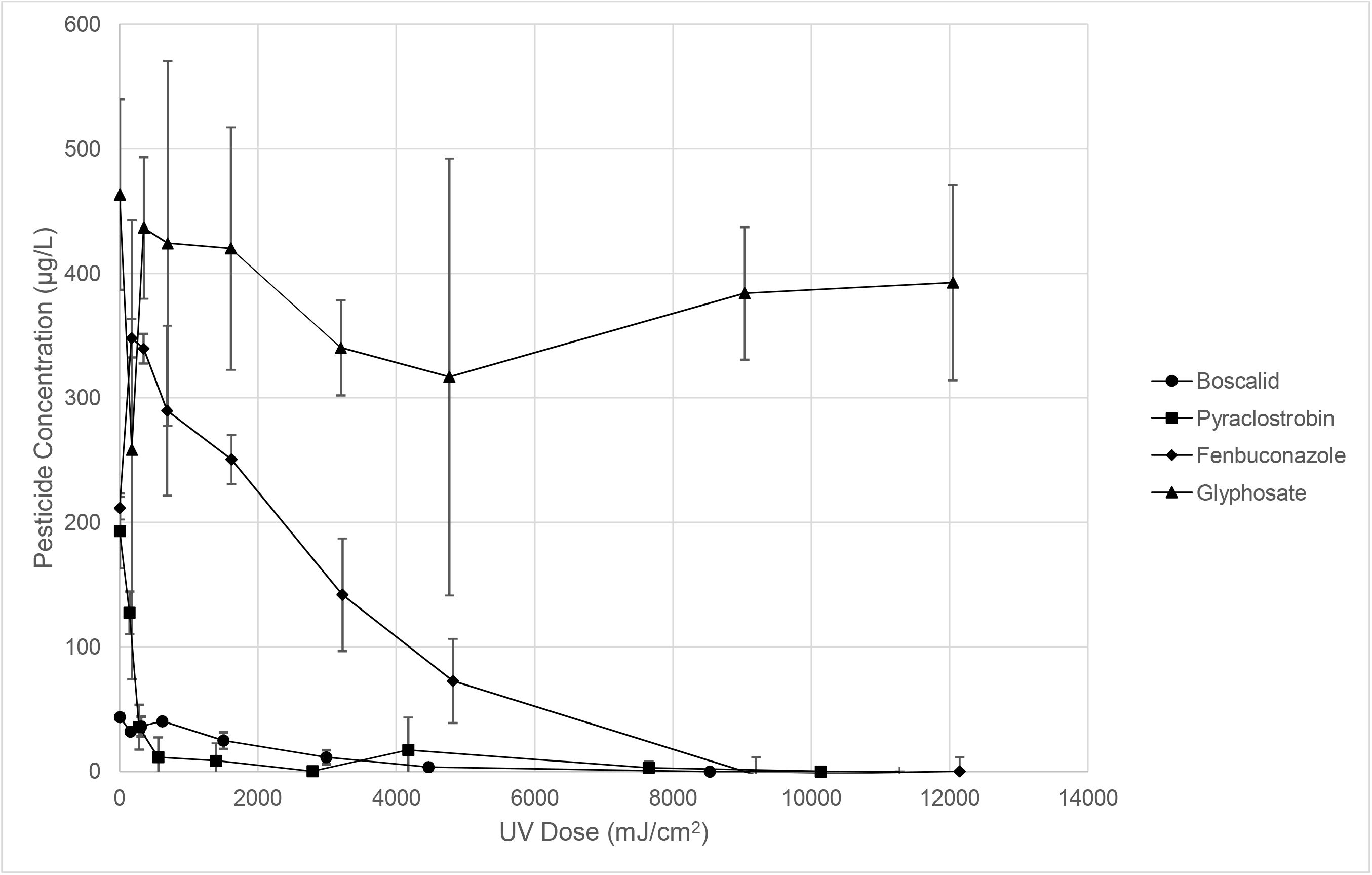
Recovered concentrations of boscalid, pyraclostrobin, fenbuconazole and glyphosate following exposure to 254 nm UV light at various UV doses on apple skin (n = 3). Error bars represent a standard deviation.

**Table 3:** Normalized recoveries of boscalid, pyraclostrobin, fenbuconazole and glyphosate after treatment on glass (top) or apple skin (bottom) with UV light at 254 nm for various times (n = 3). Data was normalized with respect to the positive controls Note that UV dose is not consistent between all samples due to variation in UV light intensity, the bracketed number is the UV dose (mW/cm^2^). Superscripts denote a significant difference between labelled exposure and ^a^ control, ^b^ 30s, ^c^ 1 min, ^d^ 2 min, ^e^ 5 min, ^f^ 10 min, ^g^ 15 min.

| Compound | Control | 30 s | 1 min | 2 min | 5 min | 10 min | 15 min | 30 min | 40 min |
| --- | --- | --- | --- | --- | --- | --- | --- | --- | --- |
| Glass |  |  |  |  |  |  |  |  |  |
| Boscalid | 1<br>(0) | 0.654<br>(150) | 0.826<br>(300) | 0.869<br>(600) | 0.775<br>(1500) | 0.697<br>(2964) | 0.747<br>(4393.5) | 0.332<sup>abcdef</sup><br>(8280) | 0.221<sup>abcdefg</sup><br>(11040) |
| Pyraclostrobin | 1<br>(0) | 0.947<sup>a</sup><br>(112.5) | 0.696<sup>ab</sup><br>(225) | 0.284<sup>abc</sup><br>(450) | 0.066<sup>abc</sup><br>(1125) | 0.005<sup>abcd</sup><br>(2257.5) | 0.009<sup>abcd</sup><br>(3396) | 0<sup>abcd</sup><br>(6399) | 0<sup>abcd</sup><br>(8490) |
| Fenbuconazole | 1<br>(0) | 1.121<br>(213.6) | 1.166<br>(422.4) | 1.105<br>(825.6) | 1.006<sup>a</sup><br>(1677) | 0.926<sup>e</sup><br>(3321) | 0.714<sup>e</sup><br>(4935) | 0.346<sup>abefg</sup><br>(9342) | 0.295<sup>aefg</sup><br>(12324) |
| Glyphosate | 1<br>(0) | 0.630<br>(175.8) | 0.785<sup>a</sup><br>(351) | 0.859<br>(699) | 0.799<br>(1650) | 0.831<br>(3273) | 0.880<br>(4860) | 0.795<sup>a</sup><br>(9126) | 1.000<br>(12060) |
| Apple |  |  |  |  |  |  |  |  |  |
| Boscalid | 1<br>(0) | 0.735<br>(153) | 0.829<br>(306.6) | 0.923<br>(613.2) | 0.569<sup>a</sup><br>(1497) | 0.263<sup>acd</sup><br>(2985) | 0.083<sup>abcde</sup><br>(4464) | 0.000<sup>abcde</sup><br>(8532) | 0<sup>abcde</sup><br>(11274) |
| Pyraclostrobin | 1<br>(0) | 0.660<sup>a</sup><br>(138.9) | 0.185<sup>ab</sup><br>(277.8) | 0.060<sup>ab</sup><br>(555.6) | 0.045<sup>ab</sup><br>(1392) | 0.001<sup>ab</sup><br>(2785.5) | 0.090<sup>ab</sup><br>(4170) | 0.016<sup>ab</sup><br>(7650) | 0.000<sup>ab</sup><br>(10134) |
| Fenbuconazole | 1<br>(0) | 1.645<sup>a</sup><br>(171) | 1.605<sup>a</sup><br>(341.7) | 1.370<br>(682.5) | 1.185<sup>b</sup><br>(1614) | 0.671<sup>bcde</sup><br>(3219) | 0.345<sup>abcde</sup><br>(4815) | 0<sup>abcdef</sup><br>(9198) | 0.001<sup>abcdef</sup><br>(12144) |
| Glyphosate | 1<br>(0) | 0.558<br>(173.4) | 0.942<br>(346.2) | 0.916<br>(690) | 0.907<br>(1605) | 0.735<br>(3195) | 0.684<br>(4764) | 0.829<br>(9036) | 0.847<br>(12048) |

**Table 4:** Molar degradation of boscalid, pyraclostrobin, fenbuconazole and glyphosate after irradiation with UV light (254 nm) for various doses on glass (n = 3).

|  | **Boscalid** |  | **Pyraclostrobin** |  | **Fenbuconazole** |  | **Glyphosate** |  |
| --- | --- | --- | --- | --- | --- | --- | --- | --- |
|  | Num. Moles degraded from control (mmol) | St Dev | Num. Moles degraded from control (mmol) | St Dev | Num. Moles degraded from control (mmol) | St Dev | Num. Moles degraded from control (mmol) | St Dev |
| **Pos Con** | 0 | ±0.013 | 0 | ±0.049 | 0 | ±0.273 | 0 | ±0.329 |
| **30 s** | 0.096 | ±0.021 | 0.076 | ±0.118 | -0.141 | ±0.199 | 1.249 | ±0.452 |
| **1 m** | 0.048 | ±0.012 | 0.437 | ±0.115 | -0.193 | ±0.195 | 0.725 | ±0.252 |
| **2 m** | 0.036 | ±0.018 | 1.030 | ±0.046 | -0.123 | ±0.194 | 0.475 | ±0.296 |
| **5 m** | 0.063 | ±0.012 | 1.343 | ±0.038 | -0.007 | ±0.194 | 0.677 | ±0.248 |
| **10 m** | 0.084 | ±0.011 | 1.431 | ±0.035 | 0.086 | ±0.230 | 0.569 | ±0.235 |
| **15 m** | 0.070 | ±0.017 | 1.424 | ±0.036 | 0.333 | ±0.197 | 0.404 | ±0.475 |
| **30 m** | 0.186 | ±0.011 | 1.441 | ±0.035 | 0.761 | ±0.196 | 0.692 | ±0.304 |
| **40 m** | 0.217 | ±0.010 | 1.438 | ±0.035 | 0.821 | ±0.194 | 0.000 | ±0.241 |

On glass (Fig. 6), boscalid was significantly degraded by the combined treatment within 1 min of exposure (279.6 mJ/cm^2^) by 0.130 ± 0.043 mmol (~56%) (Table 5). Within 10 min (2772 mJ/cm^2^) of exposure, a decrease in residue concentration of 93% was achieved and the degradation was not significantly different from the 40 min exposure (Table 5). On apple skin (Fig. 7), a significant decrease of boscalid residues took considerably longer (15 min, 4128 mJ/cm^2^) than on glass (Table 6). Complete degradation of boscalid was achieved after 30 min (8154 mJ/cm^2^) on the surface of the apple (Table 6). No previous studies of boscalid degradation using an AOP that combines UV light and hydrogen peroxide have been reported though Lagunas-Allué et al. (2010) have reported complete degradation of boscalid by UV photodegradation catalyzed by TiO_2_.

**Figure 6:**
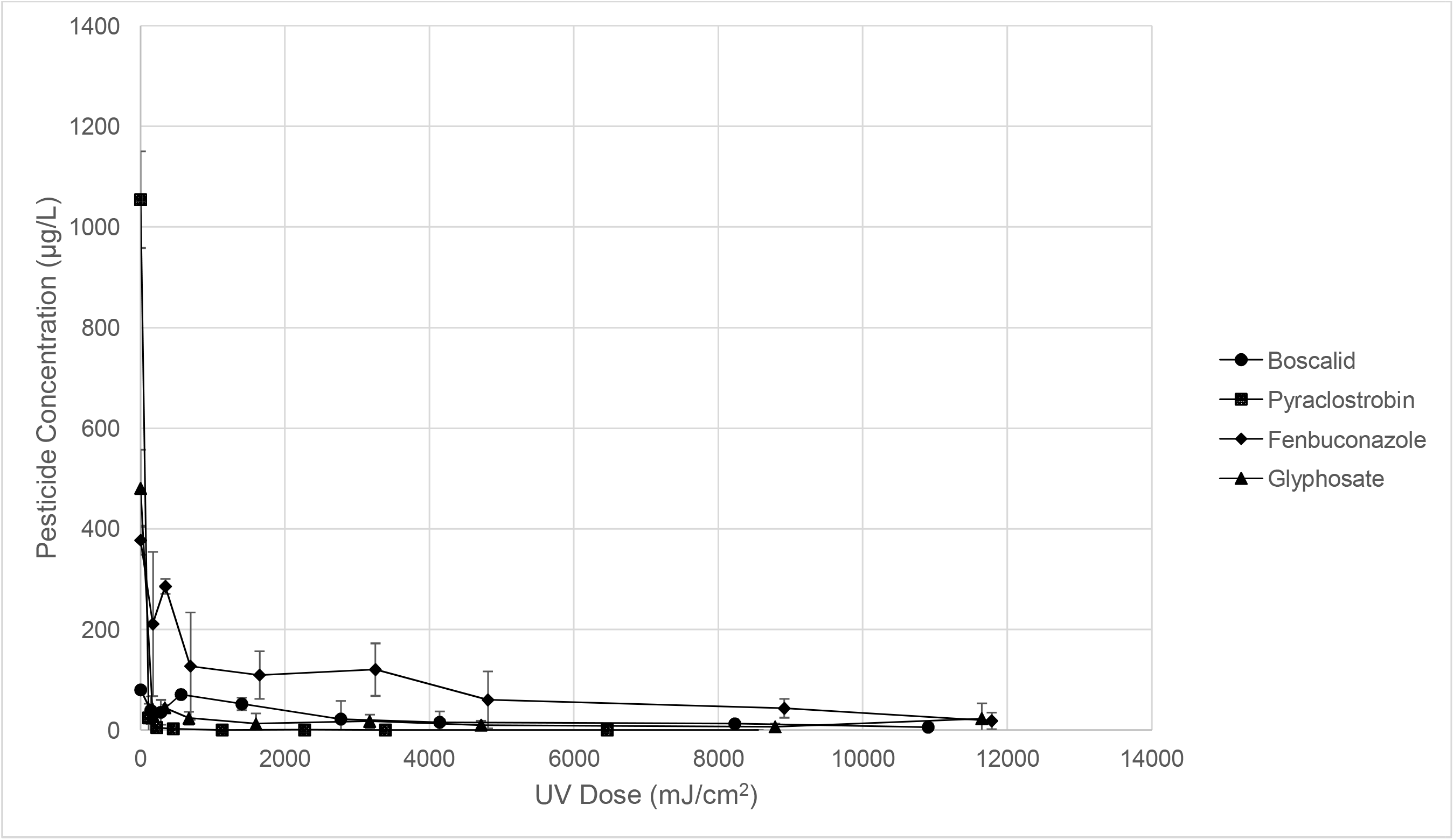
Recovered concentrations of boscalid, pyraclostrobin, fenbuconazole and glyphosate after exposure to various UV doses (254 nm) and 3% w/v hydrogen peroxide on glass (n = 3). The hydrogen peroxide was applied at 60□C for boscalid and pyraclostrobin or room temperature (23□C ± 3□C) for fenbuconazole and glyphosate. Error bars represent a standard deviation.

**Figure 7:**
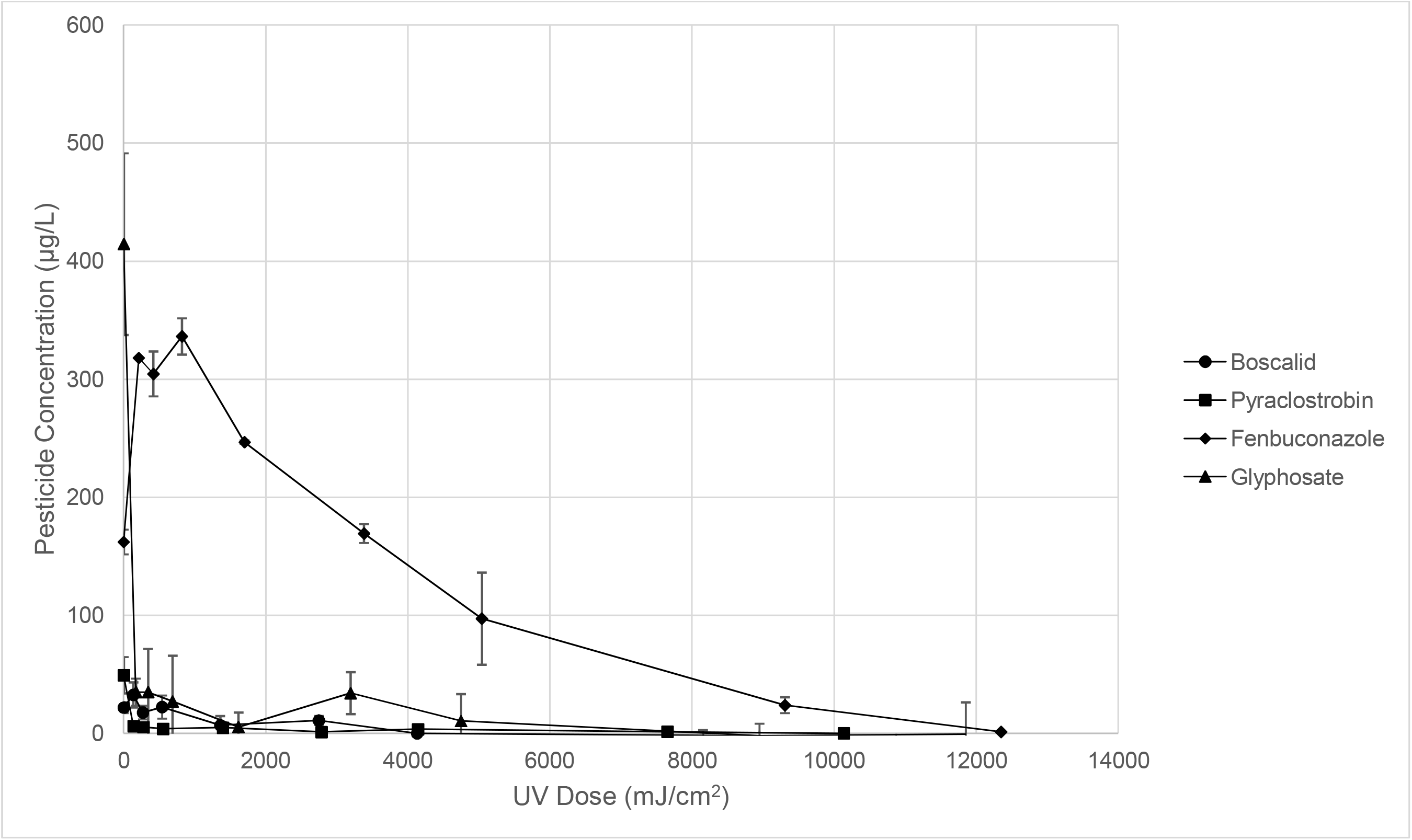
Recovered concentrations of boscalid, pyraclostrobin, fenbuconazole and glyphosate after exposure to various UV doses (254 nm) and 3% w/v hydrogen peroxide on apple skin (n = 3). The hydrogen peroxide was applied at 60□C for boscalid and pyraclostrobin or room temperature (23□C ± 3□C) for fenbuconazole and glyphosate. Error bars represent a standard deviation.

**Table 5:** Normalized recoveries of boscalid, pyraclostrobin, fenbuconazole and glyphosate after treatment on glass (top) or apple skin (bottom) with UV light at 254 nm and 3% w/v hydrogen peroxide for various times (n = 3). Hydrogen peroxide was applied at 60□C for boscalid and pyraclostrobin and room temperature (23 ± 3 □C) for fenbuconazole and glyphosate. Data was normalized with respect to the positive controls. Note that UV dose is not consistent between all samples due to variation in UV light intensity, the bracketed number is the UV dose (mW/cm^2^). Superscripts denote a significant difference between labelled exposure and ^a^ control, ^b^ 30s, ^c^ 1 min, ^d^ 2 min, ^e^ 5 min, ^f^ 10 min, ^g^ 15 min.

| Compound | Control | 30 s | 1 min | 2 min | 5 min | 10 min | 15 min | 30 min | 40 min |
| --- | --- | --- | --- | --- | --- | --- | --- | --- | --- |
| Glass |  |  |  |  |  |  |  |  |  |
| Boscalid | 1<br>(0) | 0.496<br>(139.8) | 0.443<br>(279.6) | 0.885<br>(559.2) | 0.649<br>(1401) | 0.275<sup>a</sup><br>(2772) | 0.194<sup>ad</sup><br>(4140) | 0.158<sup>ad</sup><br>(8226) | 0.074<sup>ad</sup><br>(10902) |
| Pyraclostrobin | 1<br>(0) | 0.023<sup>ab</sup><br>(112.5) | 0.005<sup>a</sup><br>(225) | 0.002<sup>a</sup><br>(450) | 0.000<sup>ab</sup><br>(1131) | 0.000<sup>ab</sup><br>(2271) | 0.000<sup>a</sup><br>(3390) | 0.000<sup>ab</sup><br>(6462) | 0<sup>a</sup><br>(8556) |
| Fenbuconazole | 1<br>(0) | 0.559<br>(171.3) | 0.757<br>(343.8) | 0.337<sup>a</sup><br>(691.2) | 0.290<sup>a</sup><br>(1647) | 0.319<sup>a</sup><br>(3249) | 0.159<sup>ac</sup><br>(4809) | 0.115<sup>ac</sup><br>(8910) | 0.049<sup>abc</sup><br>(11784) |
| Glyphosate | 1<br>(0) | 0.063<sup>a</sup><br>(167.7) | 0.092<sup>a</sup><br>(334.5) | 0.049<sup>a</sup><br>(667.5) | 0.026<sup>a</sup><br>(1596) | 0.036<sup>a</sup><br>(3171) | 0.021<sup>a</sup><br>(4713) | 0.014<sup>a</sup><br>(8784) | 0.047<sup>a</sup><br>(11646) |
| Apple |  |  |  |  |  |  |  |  |  |
| Boscalid | 1<br>(0) | 1.488<sup>a</sup><br>(134.1) | 0.811<sup>a</sup><br>(268.5) | 1.021<sup>a</sup><br>(538.5) | 0.318<sup>b</sup><br>(1359) | 0.503<sup>ab</sup><br>(2745) | 0.011<sup>ab</sup><br>(4128) | 0<sup>ab</sup><br>(8154) | 0<sup>abce</sup><br>(10872) |
| Pyraclostrobin | 1<br>(0) | 0.129<sup>a</sup><br>(138.9) | 0.109<sup>a</sup><br>(277.8) | 0.082<sup>a</sup><br>(555.6) | 0.100<sup>a</sup><br>(1395) | 0.028<sup>a</sup><br>(2781) | 0.075<sup>a</sup><br>(4140) | 0.023<sup>a</sup><br>(7650) | 0.005<sup>a</sup><br>(10134) |
| Fenbuconazole | 1<br>(0) | 1.960<sup>a</sup><br>(210.9) | 1.877<sup>a</sup><br>(416.7) | 2.073<sup>a</sup><br>(818.7) | 1.521<br>(1701) | 1.044<sup>bcd</sup><br>(3381) | 0.600<sup>bcde</sup><br>(5040) | 0.148<sup>abcdef</sup><br>(9306) | 0.008<sup>abcdefg</sup><br>(12348) |
| Glyphosate | 1<br>(0) | 0.085<sup>a</sup><br>(171) | 0.085<sup>a</sup><br>(342.3) | 0.066<sup>a</sup><br>(686.1) | 0.013<sup>a</sup><br>(1614) | 0.083<sup>a</sup><br>(3192) | 0.026<sup>a</sup><br>(4746) | 0<sup>a</sup><br>(8946) | 0<sup>a</sup><br>(11850) |

**Table 6:** Molar degradation of boscalid, pyraclostrobin, fenbuconazole and glyphosate after treatment with 3% w/v hydrogen peroxide (60□C for boscalid and glyphosate, room temperature (23±3□C) for pyraclostrobin and fenbuconazole) and various doses of UV (254 nm) (n = 3).

|  | **Boscalid** |  | **Pyraclostrobin** |  | **Fenbuconazole** |  | **Glyphosate** |  |
| --- | --- | --- | --- | --- | --- | --- | --- | --- |
|  | Num. Moles degraded from control (mmol) | St Dev | Num. Moles degraded from control (mmol) | St Dev | Num. Moles degraded from control (mmol) | St Dev | Num. Moles degraded from control (mmol) | St Dev |
| **Pos Con** | 0 | ±0.018 | 0 | ±0.203 | 0 | ±0.042 | 0 | ±0.030 |
| **30 s** | 0.118 | ±0.048 | 2.658 | ±0.149 | 0.495 | ±0.247 | 2.665 | ±0.032 |
| **1 m** | 0.130 | ±0.043 | 2.707 | ±0.143 | 0.272 | ±0.039 | 2.583 | ±0.030 |
| **2 m** | 0.027 | ±0.014 | 2.714 | ±0.143 | 0.744 | ±0.185 | 2.704 | ±0.048 |
| **5 m** | 0.082 | ±0.024 | 2.719 | ±0.143 | 0.796 | ±0.086 | 2.770 | ±0.073 |
| **10 m** | 0.170 | ±0.062 | 2.719 | ±0.143 | 0.764 | ±0.094 | 2.742 | ±0.050 |
| **15 m** | 0.188 | ±0.039 | 2.720 | ±0.143 | 0.943 | ±0.101 | 2.786 | ±0.036 |
| **30 m** | 0.196 | ±0.014 | 2.719 | ±0.143 | 0.992 | ±0.044 | 2.805 | ±0.031 |
| **40 m** | 0.216 | ±0.019 | 2.720 | ±0.143 | 1.066 | ±0.041 | 2.709 | ±0.106 |

Please note that while there has been a focus on comparable studies of oxidative degradation of the pesticides used in the current study there are many other methods that could be additionally compared such as washing, environmental processes, pulsed light etc.

### 3.2 Pyraclostrobin

Pyraclostrobin residues were not significantly decreased at any of the treatment temperatures on glass or apple skin when exposed to 3% hydrogen peroxide (Tables 1 & 2). While degradation at each of the 3% hydrogen peroxide application temperatures was not significantly different from the control, there was a general trend of increased molar degradation with increased temperature, and so 60□C was selected as the treatment temperature for the AOP owing to the greater molar decrease at that temperature (0.055 ± 0.031 mmol ~7% on glass, 0.086 ± 0.032 mmol ~23% on apple) (Tables 1 & 2). Other studies have observed that oxidation can cause a decrease of pyraclostrobin residues, but none have attempted to investigate oxidative degradation on the surface of fruit (Liu & Jiang, 2017, Zhao & Kong, 2018). Zhao & Kong (2018) examined the elimination of pyraclostrobin by electro-Fenton reaction (abiotic reactor), microbial fuel cell (reactor with disconnected electrodes) and a paired microbial fuel cell and electro-Fenton reactor. After letting the bacteria equilibrate for 6 months, 30 mg/L of pyraclostrobin was added to each reactor and decrease in residues was observed for 120 h following. While there was no significant elimination of pyraclostrobin in the abiotic reactor there was 75% degradation by microbial fuel cell alone and 100% degradation was observed in the paired reactor (concentration was below the limit of detection) (Zhao & Kong, 2018). Elimination of pyraclostrobin was therefore primarily attributed to bacterial degradation. Conversely to the elimination of pyraclostrobin observed in this study and by Zhao & Kong (2018), pyraclostrobin has been shown to be effectively degraded in water by more than 90% with hydrogen peroxide as the oxidant within 12 h when catalyzed by coke and nitric acid (Liu & Jiang, 2017). The timescale for pyraclostrobin degradation in these studies was much greater than that used for the current study. Additionally, while degradation by hydrogen peroxide was not catalyzed in this study Zhao and Kong (2018) added additional energy to the reactor cell in the form of electricity and Liu and Jiang (2017) catalyzed the reaction with coke and nitric acid.

Pyraclostrobin was the most affected by UV irradiation among the pesticides tested with a nearly 100% decrease in residue concentration (2.658 ± 0.149 mmol) within 30 s (112.5 mJ/cm^2^) on glass and 2 min (555.6 mJ/cm^2^) on apple skin (Tables 3 & 4). Zeng et al. (2019) have previously studied the photolytic and hydrolytic degradation of pyraclostrobin in aqueous solutions under UV lamp and sunlight. While the UV dose from the lamp was not reported, the sunlight averaged 840 mW/cm^2^, much higher than the incident light from the lamps used in this study (5.01 ± 0.66 mW/cm^2^). A half-life of 2.96-h and 3.69-h was reported for pyraclostrobin in paddy water under irradiation by the lamp and sunlight, respectively, showing that while sunlight was effective at degrading pyraclostrobin residues, the efficacy is diminished compared to that observed in the present study (Zeng et al., 2019). Lagunas-Allué et al. (2012) have reported significant degradation of pyraclostrobin after irradiation at 365 nm of 25 mL of a 2.3 mg/L aqueous solution of pyraclostrobin which suggests pyraclostrobin may be susceptible to a wider range of UV light than boscalid. Different transformation products of pyraclostrobin were identified by Zeng et al. (2019) and Lagunas-Allué et al. (2012) which suggests that the chromophore experiencing photoelectron transfer mediated transformation may differ depending on the wavelength of incident UV light. It should be noted that Lagunas-Allué et al. (2012) studied the transformation products of the TiO_2_ photocatalyzed elimination of pyraclostrobin which could also explain the difference in the observed oxidation products compared to Zeng et al. (2019).

Similar to UV exposure, pyraclostrobin was highly susceptible to the AOP on both glass and apple skin (Tables 5 & 6). Exposures on glass resulted in ~98% (2.658 ± 0.149 mmol) degradation within 30 s of exposure (112.5 mJ/cm^2^) and >99.9% degradation occurred within 5 min (1131 mJ/cm^2^) of exposure (Table 5). Similarly, pyraclostrobin residues were significantly degraded within 30 s (138.9 mJ/cm^2^) of exposure on apple skin, though degradation was lower at ~87% than that observed on glass (Table 5). After 10 min (2781 mJ/cm^2^) of exposure ~99% of pyraclostrobin was degraded, which was not significantly different from the degradation observed after 40 min (10134 mJ/cm^2^) of exposure (Table 5). No previous studies of pyraclostrobin degradation using an AOP combining UV light and hydrogen peroxide have been reported in the literature but Lagunas-Allué et al.(2012) have reported that exposure to UV can result in complete elimination of pyraclostrobin within 60 min when catalyzed by TiO_2_.

### 3.3 Fenbuconazole

Fenbuconazole degradation was not statistically significant on glass or apple skin by 3% hydrogen peroxide at any tested temperature though, as with boscalid on apple skin, there was a significant difference in the amount of fenbuconazole recovered at 30□C, 40□C and 50□C compared to the control (Tables 1 & 2). Unlike the slight molar degradation trend observed for pyraclostrobin with increasing temperature there was no such trend observed for fenbuconazole; therefore, room temperature was decided upon as the appropriate application temperature. No studies examining the degradation of fenbuconazole by hydrogen peroxide were found in the literature to date.

Fenbuconazole was more stable under UV irradiation than the other fungicides tested (Figs. 4 & 5). On the glass and apple skin, it took 30 min for the degradation of ~70% (0.821 ± 0.194 mmol) and 100% of fenbuconazole respectively (Tables 3 & 4). Statistically significant degradation was faster on apple skin (15 min, 4815 mJ/cm^2^) than on glass (30 min, 9342 mJ/cm^2^) (Tables 3 & 4). Lassalle et al. (2015) had previously shown cyclization of fenbuconazole as a result of irradiation with UV light of an aqueous solution in 120 mL quartz tubes in a sonicator using a high-pressure mercury lamp producing light between 200 nm and 650 nm. The degradation of fenbuconazole was not quantified by Lassalle et al. (2015) though formation and identification of eight isomers of fenbuconazole suggests there was a reduction in the amount of fenbuconazole in the solution.

Fenbuconazole was more resistant to the AOP than the other pesticides, which is not unexpected considering the decreased individual efficacy of hydrogen peroxide and UV light on fenbuconazole elimination compared to other pesticides (Tables 5 & 6). On glass, fenbuconazole was significantly degraded by 24% (0.272 ± 0.039 mmol) after an exposure of 1 min (343.8 mJ/cm^2^) (Table 5). This decrease was considerably less than those observed for the other pesticides when treated on glass (Tables 5 & 6). After 15 min (4809 mJ/cm^2^) of exposure, degradation of fenbuconazole was ~95% (1.066 ± 0.041 mmol) which was not significantly different from the 40 min (11784 mJ/cm^2^) exposure (Table 5). Fenbuconazole was even more resistant to the AOP treatment when applied on apple skin (Tables 5 & 6). Degradation was only significant from the control after 30 min (9306 mJ/cm^2^) of exposure which corresponded with a decrease in residue of ~85% (Table 5). The residue degradation reached ~99% after 40 min (12348 mJ/cm^2^) of exposure (Tables 5 & 6). This is the first study to investigate the potential degradation of fenbuconazole by an AOP.

### 3.4 Glyphosate

Glyphosate was the residue that was most degraded by 3% w/v hydrogen peroxide. On glass and apple, all tested temperatures resulted in significant residue degradation ranging from 3.060 ± 0.093 mmol to 2.757 ± 0.206 mmol (~96% and ~87% respectively) at 60□C and 40□C on glass, respectively (Tables 1 & 2). On apple skin, the degradation was as high as ~98% at 30□C compared to ~69%, ~66% and ~63% at 40□C, 50□C and 60□C respectively (Tables 1 & 2). While there was no significant difference between treatment temperatures on glass, there was greater degradation observed on apple skin at lower temperatures (Fig. 3). Therefore, room temperature was selected as the optimal application temperature for glyphosate. The elimination of glyphosate due to hydrogen peroxide observed in this study is contradictory to that observed by Junges et al. (2013) and Manassero et al. (2010) where no significant degradation of glyphosate was observed due to hydrogen peroxide alone in an aqueous solution. While in the current study there was relatively more glyphosate (2.957 mmol) compared to hydrogen peroxide (0.088mmol) the opposite was the case for Manassero et al. (0.589 mmol and 4.412 mmol of glyphosate and hydrogen peroxide respectively) (2010) and Junges et al. (0.737 mmol and 7.353 mmol of glyphosate and hydrogen peroxide respectively) (2013). Both studies used similar apparatuses in their studies. Glyphosate and hydrogen peroxide were added to water in a storage container where it was pumped through an annular reactor containing ultraviolet lamps. The reactors had total volumes of 2000 cm^3^ and ~2500 cm^3^ in the studies by Manassero et al. (2010) and Junges et al. (2013), respectively. In preliminary tests Manassero et al. (2010) exposed 0.589 mmol of glyphosate with 4.412 mmol of hydrogen peroxide for 3-h without UV irradiation and observed no change in glyphosate concentration. Junges et al. (2013) used a similar 0.737 mmol of glyphosate and 7.353 mmol of hydrogen peroxide and also ran the reactor in the absence of UV irradiation observing no noticeable change in glyphosate concentration after 4-h. While the concentration of hydrogen peroxide used by Manassero et al. (2010) and Junges et al. (2013) is likely to quench the formation of the hydroxyl radical compared to the concentration used in the current study, it is unlikely that the treatment conditions in any of the three studies would drive hydroxyl radical formation. It is more likely that the method of hydrogen peroxide application in this study enabled the applied glyphosate residue to slightly acidify the 3% hydrogen peroxide solution driving formation of hydrogen peroxide’s conjugate base and thus an acid-base reaction with glyphosate (discussed further in section 3.5). The volume of water used in the reactors by Manassero et al. (2010) and Junges et al. (2013) would not permit a similar acidification to occur and may explain the lack of glyphosate elimination observed in those studies.

Degradation of glyphosate residue was not statistically significant when exposed to UV light after exposure to 12,060 mJ/cm^2^ or 12,048 mJ/cm^2^ (40 min of irradiation) on glass and apple skin, respectively (Figs. 4 & 5). The 1 min (351 mJ/cm^2^) exposure on glass was an anomaly as it was significantly different from the control with an ~21% degradation (1.249 ± 0.452 mmol) (Tables 3 & 4). The decreased concentration in the 1 min treatment was likely due to poor recovery of the glyphosate from the glass as the observed degradation was greater in this exposure than for longer exposures (greater UV dose). Glyphosate’s resistance to photodegradation has been reported by Rueppel et al. (1977) and further confirmed by Manassero et al. (2010) and Junges et al. (2013). Use of a Crosby reactor for a 48-h exposure (equivalent to sixteen 8-h days sunlight (Rueppel et al., 1977)), and irradiation of 50 mg/L aqueous solutions of glyphosate by a 40 W Heraeus UV lamp for up to 4-h (Junges et al., 2013; Manassero et al., 2010) both failed to result in significant residue degradation. The resistance of glyphosate to UV exposure is likely due to the relative lack of chromophores compared to the fungicides (Manassero et al., 2010).

Similar to pyraclostrobin, glyphosate was highly susceptible to the AOP treatment on both glass and apple skin (Figs. 6 & 7). On both substrates, residue degradation was significant within 30 s (167.7 mJ/cm^2^ and 171 mJ/cm^2^ for glass and apple skin, respectively) of exposure with ~94% (2.665 ± 0.032 mmol) and ~91% of glyphosate being degraded from glass and apple skin, respectively (Tables 5 & 6). After only 30 s of exposure, degradation was not statistically different from the 40 min (11646 mJ/cm^2^ and 11850 mJ/cm^2^ for glass and apple skin respectively) exposure; achieving a decrease of ~95% and ~100% for glass and apple skin, respectively (Table 5). AOP exposures with UV and hydrogen peroxide have previously been shown to be effective at significantly decreasing glyphosate residues (Junges et al., 2013; Manassero et al., 2010). Junges et al. (2013) and Manassero et al. (2010) studied residue degradation of a solution of glyphosate (50 mg/L) in an annular reactor using 120 mg/L of hydrogen peroxide (note that Manassero et al.(2010) varied this condition) and a germicidal lamp (254 nm). Under optimal conditions Junges et al. (2013) reported 90% degradation of glyphosate after 6 h of exposure while Manassero et al. (2010) only observed a 63.5% degradation after 5 h. The decrease observed in the current study occurred much more rapidly than that reported by Junges et al. (2013) and Manassero et al. (2010) This is likely due to the much higher concentration of hydrogen peroxide used in those studies resulting in quenching of production of the hydroxyl radical.

### 3.5 Comparison of Treatments

The degree of pesticide residue degradation among the optimal temperature of hydrogen peroxide, longest exposure durations for UV light and the longest exposure duration of the AOP were compared to identify any significant differences (Table 7). For the fungicides, UV light and the AOP were the most effective methods of degradation with hydrogen peroxide being significantly less effective for all three. Glyphosate was most effectively degraded by treatment with hydrogen peroxide and the AOP, significantly more than the degradation observed by UV light alone. While there appears to be a difference between residue degradation by UV light and the AOP for boscalid and fenbuconazole, the difference is not statistically significant. Thus, there is not a statistically significant difference in residue degradation with the AOP compared to the optimal individual UV light or hydrogen peroxide treatments for either of these pesticides.

**Table 7:** Comparison between hydrogen peroxide, UV light and the AOP for boscalid, pyraclostrobin, fenbuconazole and glyphosate. The displayed percentages represent the % degradation based on molar degradation between the positive control and longest treatment time (for UV and AOP treatments) or optimal hydrogen peroxide temperature (60□C for boscalid and glyphosate, room temperature (23±3□C) for pyraclostrobin and fenbuconazole) on glass.

|  | **Boscalid** |  | **Pyraclostrobin** |  | **Fenbuconazole** |  | **Glyphosate** |  |
| --- | --- | --- | --- | --- | --- | --- | --- | --- |
|  | % Degraded | St Dev | % Degraded | St Dev | % Degraded | St Dev | % Degraded | St Dev |
| **H<sub>2</sub>O<sub>2</sub>** | 38.38 | ±3.00 | 3.34 | ±1.92 | 3.12 | ±2.10 | 96.04 | ±4.78 |
| **UV Light** | 77.94 | ±5.79 | 100.03 | ±4.84 | 70.49 | ±26.19 | 0.00 | - |
| **AOP** | 92.63 | ±11.96 | 100.00 | ±10.54 | 95.07 | ±5.70 | 95.26 | ±3.91 |

While there is no difference in the total decrease in residue concentration, there is a significant difference in the amount of time required to reach the maximum level of degradation using UV light or the AOP. Under UV light alone, maximum degradation was achieved after 40 min, 10 min and 30 min for boscalid, pyraclostrobin and fenbuconazole, respectively, but that was reduced to 10 min, 30 s, and 15 min under the AOP (Tables 3 & 5). Due to a single treatment duration being tested with hydrogen peroxide, a similar comparison for glyphosate is not possible. However, given that the increased rate of residue degradation for the fungicides when exposed to AOP is likely due to production of the hydroxyl radical owing to photodegradation of hydrogen peroxide by the UV light, it is likely that glyphosate would similarly be degraded more quickly by the AOP than by hydrogen peroxide alone.

Differences in the degradation of the pesticides compared to one another and between treatments is due to differences in their structure. Elimination by UV light requires the presence of a chromophore-a structure capable of absorbing light of a specific wavelength. Benzene is a common structural element among the fungicides in this study and is capable of absorbing light up to 200 nm. The presence of adjacent atoms and functional groups can shift the wavelength of light absorbed, and so the fungicides can absorb the 254 nm UV light used in the current study while glyphosate cannot. The energy gained by absorption of a photon can be re-emitted or result in electron transfer between the fungicide and the surrounding matrix, either the apple epicuticular waxes or the air (Zhang, 1997; Hasegawa, 2006). In the case of energy transfer to the air, there is the potential for formation of reactive oxygen species that would drive transformation of the pesticide residues and result in the degradation observed in this study (Zhang, 1997; Hasegawa, 2006). Reactivity with hydrogen peroxide differs. At a neutral pH, glyphosate has three negatively charged functional groups (pKa <2, 2.6 and 5.6) and is therefore acidic. Hydrogen peroxide itself is a weak acid (pKa 11.7). Following addition of the hydrogen peroxide droplet to a glyphosate treated surface may sufficiently acidify the solution to drive production of the conjugate base of hydrogen peroxide resulting in an acid-base reaction between glyphosate and hydrogen peroxide and thus the transformation of glyphosate (into aminomethylphosphonic acid for example). As there would be little to catalyse the formation of the hydroxyl radical during the hydrogen peroxide exposures, it is unlikely that glyphosate residue degradation is occurring via that method. While a detailed discussion of the reactivity of the functional groups in each of the pesticides tested is beyond the scope of the current study, it should be noted that differences in the reactivity of the functional groups, for example the amide of boscalid, cyano- or triazole of fenbuconazole, chlorobenzene common to all of the fungicides, carboxylic acids of pyraclostrobin and glyphosate or the phosphate of glyphosate, in each pesticide could also influence their reactivity under the oxidative treatments used in this study.

In general, degradation of pesticides occurred more effectively on the surface of glass than on the surface of apple skins (Tables 1, 3 & 5). This is likely due to pesticides being absorbed into the apple skin. It is also important to note that exposure of apple skins to hydrogen peroxide at different temperatures could have influenced absorption of the pesticide, and consequently, to their degradation. Changes in temperature on the surface of the apple skin could affect the epicuticular waxes on the surface of the apple skin, which could increase absorption of the pesticide residues into the skin (Kirkwood, 1999; Riccio et al, 2006). The increased absorption of pesticides into the skin of the apple could make them less susceptible to transformation by the different oxidative treatments (Angioni et al., 2004; Riccio et al., 2006), which could explain the results observed in this study.

## 4 Conclusion

Only boscalid and glyphosate were significantly degraded by hydrogen peroxide with boscalid degradation improving with greater application temperature, an opposite trend was observed for glyphosate. The fungicides boscalid, pyraclostrobin and fenbuconazole were significantly degraded by irradiation with UV light while glyphosate was not. Residue degradation by AOP was found to occur more rapidly for boscalid, pyraclostrobin and fenbuconazole than by UV light alone. Glyphosate degradation under AOP was comparable to that observed during exposure to just hydrogen peroxide. These results indicate that an AOP approach may be capable of residue degradation among a larger variety of pesticide residues than choosing a single oxidative process because certain pesticides will be more susceptible to certain oxidative processes than others.

## Acknowledgements

The authors wish to thank the Natural Science and Engineering Research Council of Canada for their support through Engage (#401429) and Discovery (#401357) grants. We also wish to thank the Ontario Centres for Excellence (VIP I #053858) and Moyer’s Apple Products for their funding support and the donation of apples and Clean Works Corporation for the laboratory scale Advanced Oxidative Process reactor.

