## Supplementary Information for "Degradation of *boscalid, pyraclostrobin, fenbuconazole*, and *glyphosate* residues by an advanced oxidative process utilizing ultraviolet light and hydrogen peroxide"

**Table S1**: Summary of previous studies examining degradation of pesticide residues. Amount degraded is provided in the same order as the methods of degradation and under optimal experimental conditions for each study. For studies with more than one pesticide the amount degraded is presented with the first number representing the first listed pesticide and so on.

| **Authors** | **Method of Pesticide Degradation** | **Pesticides Studied** | **Amount Degraded** | **Matrix from which pesticide was degraded** |
| --- | --- | --- | --- | --- |
| Crowe et al. 2006 | Aqueous washes of:  O_3_  H_2_O_2_  Cl_2_  UV  UV/H_2_O_2_  UV/Cl_2_  UV/H_2_O_2_/O_3_ | Phosmet | 57.7%  23.2%  46%  31.2%  39.4%  Not Significant  Not Significant | Surface of lowbush blueberries |
| Hwang et al. 2001a | Aqueous washes of:  Cl_2_  ClO_2_  O_3_  HPA | Mancozeb | 99.9%  87%  97%  99% | Surface of Apple |
| Hwang et al. 2001b | Aqueous:  O_3_  HPA | Mancozeb | 100%  75% | Aqueous solution |
| Hwang et al. 2002 | Aqueous washes of:  Water  Cl_2_  ClO_2_  O_3_  HPA  Also compared processing methods | Mancozeb | 48.1%  100%  87.9%  100%  82.1% | Surface of Apple |
| Hwang et al. 2003 | Aqueous washes of:  Cl_2_  O_3_  Focused on transformation pathway | Mancozeb | Degradation was not quantified | Aqueous solution |
| Cengiz and Certel, 2014 | Aqueous washes of:  Water  Cl­_2_  H_2_O_2_  O_3_ | Mancozeb | 22%  71%  65%  60% | Surface of Tomato |
| Ong et al., 1996 | Aqueous washes of:  Cl­_2_  O_3_  Water | Azinphos-methyl  Captan  Fometanate hydrochloride | 83%/77%/50%  75%/72%/46%  53%/50%/23% | Surface of Apple |
| Wu et al., 2007 | Aqueous wash of O_3_ | Diazinon  Methyl parathion  Parathion  Cypermethrin | 99%/60%/~85%/  ~87% | Aqueous solution  Wild mustard |
| Wu et al., 2009 | Aqueous wash of O_3_ | Diazinon  Methyl parathion  Parathion | Degradation was not quantified | Aqueous Solution |
| Lester et al., 2017 | Sunlight  Aqueous washes of:  O_3_  H_2_O_2_ | Chlorpyrifos | ~68%  7%  Not reported | Experimental surface  Leaves |
| Savi et al., 2016 | Gaseous O_3_ | Bifenthrin  Pirimiphos-methyl | 37.5%/71.1% | Wheat Grain |
| Savi et al., 2015 | Gaseous O­_3_ | Deltamethrin  Fenitrothion | 89.8%/66.7% | Wheat Grain |
| Kusvuran et al., 2012 | Aqueous washes of:  O_3_  Water | Chlorpyrifos ethyl  Tetradifon  Chlorothalonil | 94.2%/98.6%/100%  39.7%/31.4%/24.7% | Surface of lemon, orange, and grapefruit |
| Sadło et al., 2017 | Water  Aqueous O_3_  Gaseous O_3_ | Captan  Boscalid  Pyraclostrobin | 81%/67.3%/42.5%  ~81%/40.4%/20%  None/42.3%/32.5% | Surface of Apple |
| Hoff et al. 2019 | Aqueous O_3_  Solar Still | Epoxiconazole  Pyraclostrobin | 73%/90.8%  >99.995%/99.99% | Effluent water |
| Karaca et al., 2012 | Gaseous O_3_ | Boscalid  Iprodione  Fenhexamid  Cyprodinil  Pyrimethanil | 46.2%/23.9%/64.5%/  34.7%/51.6% | Surface of Grapes |
| Manassero et al., 2010 | Aqueous H­_2_O­_2_  UV  UV/H_2_O_2_ | Glyphosate | None  None  70% | Aqueous solution |
| Junges et al., 2013 | Aqueous H_2_O_2_  UV  UV/H_2_O_2_ | Glyphosate | None  None  70% | Aqueous solution |
| Rueppel et al., 1977 | Environmental conditions | Glyphosate | 90% | Soil  Aqueous solution |
| Zhao and Kong, 2018 | Electro-Fenton  Microbial metabolism  Both | Pyraclostrobin | 10%  76.7%  100% | Aqueous solution |
| Liu and Jiang, 2017 | H_2_O_2_ generated by coke and nitric acid | Pyraclostrobin | 100% | Aqueous solution |
| Lassalle et al., 2014 | UV | Boscalid | 90% | Aqueous solution |
| Lassalle et al., 2015 | UV | Fenbuconazole | Degradation was not quantified | Aqueous solution |
| Lagunas-Allué et al., 2010 | TiO_2_ catalyzed photolysis | Boscalid | 100% | Aqueous solution |
| Lagunas-Allué et al., 2012 | TiO_2_ catalyzed photolysis | Pyraclostrobin | 100% | Aqueous solution |
| Zeng et al., 2019 | UV  Hydrolysis | Pyraclostrobin | 100%  79.2% | Aqueous solution |
| Kaur et al., 1997 | Environmental conditions | Methyl parathion  Malathion | Degradation was not quantified | Radish  Carrot  Soil |
| Slotkin et al., 2008 | UV collimated beam | Chlorpyrifos | 70% | Aqueous solution |

**Table S2**: Recovery experiments with boscalid testing methanol, acetonitrile, and acetone as recovery solvents from apple skin segments. Spikes were 100µL aliquots of 2000 µg/L solution and were recovered with 1 mL of recovery solvent. Spikes were recovered immediately after complete evaporation of the application solvent. Negative controls (not shown) indicated no background boscalid on the apple recoverable by any tested solvent.

| **Application Solvent** | **Recovery Solvent** | **% Recovery ([Sample]/[Spike])** | **Standard Deviation** |
| --- | --- | --- | --- |
| Methanol | Methanol | 47.00 | 12.53 |
| Methanol | Acetonitrile | 55.67 | 8.62 |
| Acetonitrile | Methanol | 93.67 | 0.58 |
| Acetonitrile | Acetonitrile | 73.67 | 13.65 |
| Acetone | Methanol | 69.33 | 8.08 |
| Acetone | Acetonitrile | 69.33 | 5.15 |

|  | **Boscalid** | | **Pyraclostrobin** | | **Fenbuconazole** | | **Glyphosate** | |
| --- | --- | --- | --- | --- | --- | --- | --- | --- |
|  | **Recovery %** | **Std Dev** | **Recovery %** | **Std Dev** | **Recovery %** | **Std Dev** | **Recovery %** | **Std Dev** |
| **Glass** |  |  |  |  |  |  |  |  |
| **Dry** | 110.21 | 5.03 | 98.75 | 3.74 | 99.25 | 5.93 | - | - |
| **60 min** | 112.96 | 6.35 | 95.33 | 7.28 | 99.26 | 2.97 | 95.30 | 11.39 |
| **Apple** |  |  |  |  |  |  |  |  |
| **Dry** | 108.79 | 2.55 | 76.03 | 12.11 | 70.86 | 3.16 | 101.02 | 23.49 |
| **60 min** | 96.22 | 10.43 | 39.30 | 13.14 | - | - | 88.84 | 20.79 |

**Table S3**: Pesticide recoveries from glass and apple skin. Boscalid, pyraclostrobin and fenbuconazole were applied in acetonitrile and recovered with methanol. Glyphosate was applied in distilled water, dried in a ~50⁰C oven and recovered with distilled water.

**Table S4**: Summary of chromatographic conditions. Solvent A: HPLC grade water (0.1% formic acid). Solvent B: HPLC grade acetonitrile (0.1% formic acid).

| **Compound** | **Initial Conditions** | **Time Held** | **Gradient Type** | **Time for Gradient Change** | **Final Conditions** | **Time Held** | **Total Time** |
| --- | --- | --- | --- | --- | --- | --- | --- |
| Boscalid | 20% A | 3 min | Single Step | 13 min | 100% B | 5 min | 21 min |
| Pyraclostrobin | 60% A | - | Single Step | 15 min | 100% B | 2 min | 17 min |
| Fenbuconazole | 30% A | 5 min | Single Step | 7 min | 100% B | 5 min | 17 min |
| Glyphosate | 100% B | 2 min | Single Step | 5 min | 80% B | - | 7 min |

**Table S5**: Ionization mode and monitored transition for boscalid, pyraclostrobin, fenbuconazole and glyphosate.

| **Compound** | **Ionization mode** | **Monitored Transition (m/z)** |
| --- | --- | --- |
| Boscalid | +ve | 343-307 |
| Pyraclostrobin | +ve | 388-194 |
| Fenbuconazole | +ve | 337-125 |
| Glyphosate | -ve | 168-150 |

**Table S6**: Recovered concentrations of boscalid, pyraclostrobin, fenbuconazole and glyphosate after exposure to 3% hydrogen peroxide at various temperatures for 15 min on glass and apple skin (n = 3).

|  | **Control** | | **30⁰C** | | **40⁰C** | | **50⁰C** | | **60⁰C** | |
| --- | --- | --- | --- | --- | --- | --- | --- | --- | --- | --- |
|  | **Concentration (µg/L)** | **Std Dev** | **Concentration (µg/L)** | **Std Dev** | **Concentration (µg/L)** | **Std Dev** | **Concentration (µg/L)** | **Std Dev** | **Concentration (µg/L)** | **Std Dev** |
| **Glass** |  |  |  |  |  |  |  |  |  |  |
| **Boscalid** | 94.45 | 3.90 | 91.56 | 7.34 | 79.37 | 5.17 | 76.67 | 4.10 | 58.20 | 1.47 |
| **Pyraclostrobin** | 303.52 | 15.99 | 313.65 | 14.86 | 310.50 | 14.90 | 312.43 | 6.78 | 282.37 | 13.51 |
| **Fenbuconazole** | 373.54 | 20.14 | 385.20 | 16.35 | 366.35 | 10.82 | 381.20 | 21.17 | 409.23 | 36.32 |
| **Glyphosate** | 538.63 | 21.17 | <LOQ | - | 72.47 | 56.63 | 46.39 | 20.63 | <LOQ | - |
| **Apple** |  |  |  |  |  |  |  |  |  |  |
| **Boscalid** | 15.04 | 8.16 | 43.06 | 7.58 | 58.23 | 6.86 | 39.29 | 15.96 | 33.11 | 9.64 |
| **Pyraclostrobin** | 143.61 | 10.37 | 167.23 | 21.76 | 179.80 | 30.29 | 146.69 | 9.84 | 110.08 | 19.10 |
| **Fenbuconazole** | 95.58 | 39.97 | 249.31 | 15.44 | 199.80 | 22.41 | 154.80 | 18.77 | 115.73 | 0.65 |
| **Glyphosate** | 453.84 | 33.79 | < LOD | - | 141.34 | 46.35 | 155.55 | 30.10 | 166.18 | 65.75 |

**Table S7**: Recovered concentrations of boscalid, pyraclostrobin, fenbuconazole and glyphosate after irradiation with UV-C (254 nm) light after various times on glass and apple skin (n = 3).

|  | **Control** | **30 s** | **1 min** | **2 min** | **5 min** | **10 min** | **15 min** | **30 min** | **40 min** |
| --- | --- | --- | --- | --- | --- | --- | --- | --- | --- |
|  | **Conc. (µg/L)** | **Conc. (µg/L)** | **Conc. (µg/L)** | **Conc. (µg/L)** | **Conc. (µg/L)** | **Conc. (µg/L)** | **Conc. (µg/L)** | **Conc. (µg/L)** | **Conc. (µg/L)** |
| **Glass** |  |  |  |  |  |  |  |  |  |
| **Bosc.** | 95.55  ±5.60 | 62.45  ±11.41 | 78.92  ±4.84 | 83.07  ±9.22 | 74.07  ±4.00 | 66.64  ±3.14 | 71.37  ±8.52 | 31.69  ±3.20 | 21.08  ±1.82 |
| **Pyra.** | 557.53  ±23.35 | 528.02  ±75.56 | 388.24 ±73.40 | 158.17  ±20.07 | 36.60  ±10.31 | 2.59  ±3.66 | 5.11  ±6.46 | < LOD | <LOD |
| **Fenb.** | 392.33 ±112.56 | 439.68  ±27.33 | 457.30 ±16.95 | 433.66  ±10.57 | 394.66  ±9.07 | 363.22 ±72.70 | 280.20  ±22.05 | 135.91 ±19.16 | 115.76 ±11.03 |
| **Glyp.** | 569.83  ±68.10 | 358.73 ±113.44 | 447.23 ±28.26 | 489.53  ±53.40 | 455.42 ±25.58 | 473.56  ±9.35 | 501.54 ±121.38 | 452.90 ±57.17 | 569.83 ±18.64 |
| **Apple** |  |  |  |  |  |  |  |  |  |
| **Bosc.** | 43.73  ±3.64 | 32.12  ±0.26 | 36.26  ±7.88 | 40.38  ±0.50 | 24.89  ±6.78 | 11.51  ±5.66 | 3.61  ±2.38 | < LOD | < LOD |
| **Pyra.** | 193.30  ±28.27 | 127.54  ±17.18 | 35.69  ±18.00 | 11.55  ±15.87 | 8.68  ±14.13 | < LOD | 17.37  ±26.11 | 3.07  ±5.33 | < LOD |
| **Fenb.** | 211.57  ±9.21 | 348.09  ±15.54 | 339.66 ±11.85 | 289.87  ±68.24 | 250.76 ±19.67 | 142.02 ±45.24 | 72.89  ±33.83 | < LOD | < LOD |
| **Glyp.** | 463.34  ±91.68 | 258.46 ±184.29 | 436.60 ±56.86 | 424.20 ±146.62 | 420.08 ±97.37 | 340.33 ±38.25 | 316.94 ±175.43 | 384.07 ±53.25 | 392.59 ±78.36 |

**Table S8**: Recovered concentrations of boscalid, pyraclostrobin, fenbuconazole and glyphosate after AOP treatment on glass and apple skin for various durations (n = 3).

|  | **Control** | **30 s** | **1 min** | **2 min** | **5 min** | **10 min** | **15 min** | **30 min** | **40 min** |
| --- | --- | --- | --- | --- | --- | --- | --- | --- | --- |
|  | **Conc. (µg/L)** | **Conc. (µg/L)** | **Conc. (µg/L)** | **Conc. (µg/L)** | **Conc. (µg/L)** | **Conc. (µg/L)** | **Conc. (µg/L)** | **Conc. (µg/L)** | **Conc. (µg/L)** |
| **Glass** |  |  |  |  |  |  |  |  |  |
| **Bosc.** | 160.60  ±4.49 | 39.67 ±27.27 | 35.49 ±24.63 | 70.81  ±2.76 | 51.97 ±12.39 | 22.00 ±35.94 | 15.55 ±21.68 | 12.64  ±2.97 | 5.90  ±8.25 |
| **Pyra.** | 527.44 ±96.29 | 24.04 ±28.19 | 5.13  ±2.23 | 2.37  ±0.74 | < LOD | < LOD | < LOD | < LOD | < LOD |
| **Fenb.** | 949.54 ±28.80 | 210.88 ±143.31 | 285.87 ±14.78 | 127.15 ±106.70 | 109.49 ±47.28 | 120.34 ±52.15 | 59.87 ±56.54 | 43.42 ±18.62 | 18.62 ±16.44 |
| **Glyp.** | 610.75 ±76.93 | <LOQ | 44.16  ±6.46 | <LOQ | <LOD | <LOQ | <LOD | <LOD | <LOQ |
| **Apple** |  |  |  |  |  |  |  |  |  |
| **Bosc.** | 106.60  ±4.49 | 32.75 ±10.71 | 17.84  ±5.84 | 22.48  ±9.78 | 6.99  ±7.87 | 11.08  ±3.74 | < LOD | < LOD | < LOD |
| **Pyra.** | 49.48  ±15.35 | 6.37  ±0.70 | 5.38  ±1.63 | 4.04  ±1.65 | 4.94  ±2.03 | 1.37  ±1.18 | 3.69 ±0.89 | 1.48  ±2.02 | < LOD |
| **Fenb.** | 484.06 ±10.43 | 318.11 | 304.62 ±18.96 | 336.40 ±15.32 | 246.90 ±1.31 | 169.39 ±8.00 | 97.44 ±39.01 | 24.08  ±6.71 | 1.35  ±0.57 |
| **Glyp.** | 610.75 ±76.93 | 35.04 ±11.47 | 35.06 ±36.70 | <LOQ | <LOD | 34.25 ±17.71 | <LOQ | < LOD | < LOD |
